# A meta-analysis on the effects of pollutants on the earthworm microbiota

**DOI:** 10.64898/2026.09.14.751390

**Authors:** Tina M. Raassina, Linyang Sun, Anne Duplouy, Helen R. P. Phillips

## Abstract

Like for many other organisms, earthworms host a microbiota consisting of bacteria, fungi and other micro-organisms originating from their surrounding environment. In earthworms, these microbes, either associated with the gut, skin, kidney-like nephridia, gizzard or cocoons, aid their hosts with the digestion of organic matter, immune responses against pathogens, and/or detoxification. Unfortunately, pollution is an increasing threat to soil health and soil biodiversity, and numerous articles have characterized both the microbial communities associated with earthworms, as well as the effects of soil pollution on these microbial communities. Here, we aim to synthesize data from across 159 articles to identify consistent global patterns and gaps in the study of earthworms’ microbial communities. Additionally, we combined 16S rRNA metabarcoding data from almost 30 articles in a meta-analysis to evaluate the changes in the diversity of earthworm’s and surrounding soil’s bacterial microbiota after pollutant exposure. Through an evidence map, we clearly showed that most of the earthworm microbiota data came from the *Eisenia fetida* species reared in laboratory conditions, and only a handful of studies sequenced fungi (ITS) and protists (18S rRNA) in addition to bacteria and archaea (16S rRNA). The scarcity of studies from wild populations, environmental conditions, and the low microbial diversity targets and sources, cannot support a comprehensive view on earthworm’s microbial ecology. Nonetheless, results of the meta-analysis indicate that the data did not support the occurrence of significant changes in the microbiota richness and diversity as result of pollution exposure. However, the microbial communities altered their composition, becoming more dissimilar. We suspect the microbiota response to pollution to be host-species specific, but the lack of data severely limited our ability to further elucidate the effect of pollution on earthworm microbiota.

## Introduction

The role of earthworms in the provision of ecosystem functions and services, including soil-mixing, water permeability, and habitat creation, amongst others, is well documented^1–3^. The microbiota of these terrestrial organisms plays an important role in their host functioning, as well as the wider ecosystem functions earthworms provide. For example, following the ingestion of soil, earthworms can digest the soil organic matter by utilizing their own enzymes, as well as the enzymes excreted by the diverse microbes colonizing their digestive system^4,5^. Additionally, the earthworm gut microbiota is colonized by functional bacterial groups such as ammonia oxidizers, sulphate and nitrite reducers, dehalogenators^6^, and bacteria that are linked to carbon metabolism, or carbon cycling. Earthworms excrete wormcasts, whose rich-microbe composition is different to that found in the soil^7,8^. By enriching the diversity and abundance of the soil with their own microbial set^9,10^, earthworms and their wormcasts have been shown to increase plant yield, reduce soil pathogens, and thus enhance ecosystems resilience against stressors^9^, and overall health of these ecosystems.

Typically, the relative abundances of the different microbial taxa significantly differ between earthworm host species^6,11^. And although the gut microbiota of earthworms has been extensively studied^9^, their entire holobiont also include the microbes associated with their nephridia, skin, casts^9^, gizzard^12^ and cocoons^13^. Expectedly, earthworms acquire their associated microbes through ingestion of soil^14,15^, but studies have shown that their gut microbiota typically consists of only a subset of the available soil microbial species^15^.

The majority of pollutants in the soil have anthropogenic origins, for example from industrial activities, mining, waste management, and agricultural practices^16^, which are increasing with human population density^17^. As soils act like giant filters, trapping all the contaminants that are washed through the soil aggregates^18^, pollution has become one of the biggest threats to soil biodiversity and health^19^. Earlier meta-analyses have explored the global impact of pollution on earthworm species as whole organisms and have shown adverse effects of pollutants on earthworms by reducing their growth, survival^20,21^, reproduction^20^, suppressing their metabolism^22^, and decreasing communities-level abundance and diversity^19^.

Although exposure to pollutants has also been shown to affect the earthworm microbiota, results from different studies are not always consistent. For example, low-density polyethylene (LDPE) plastic has been shown to increase^23^, decrease^24^, or to have no significant effect on earthworm-associated microbial diversity^25,26^. These contrasting results could be explained by sampling differences between studies, as different earthworm species appear to respond differently to pollutants. Pollution with polyethylene (PE) plastic was found to decrease microbial diversity in *Eisenia nordenskioldi*^27^ but not in *Eudrilus eugeniae*^28^. In contrast, while the earthworm gut microbiota is highly sensitive to pollution by the heavy metal antimony (Sb)^29^, the microbiota of earthworm epidermis and cast are rather resistant to biochar-derived dissolved and particulate matter exposure^30^.

As there is no meta-analysis on the effects of pollutants on the microbiota of earthworms across species and pollutant groups, we aimed to fill this gap. Given the importance of the microbiota not only for the general biology of the species but also for the provision of ecosystem functions and healthy soils, such a general overview of the field would bring some clarity of the state of the research field on pollutant impacts on the earthworm microbiota. For the meta-analysis, articles were collected through a systematic literature search. An evidence map from the included articles was created to get an overview of studies that exist on the topic, as well as provide a roadmap for future research. Additionally, we extracted and analysed the articles associated 16S rRNA metabarcoding data to investigate any possible shifts in the community composition, diversity and richness of the earthworm microbiota after pollutant exposure. Although pollutants have shown varying effects on the earthworm microbiota depending on the pollutant and earthworm species tested, we expected pollutants to generally reduce microbial diversity and richness, as well as shift their community composition to become more dissimilar given the importance of the microbiota on earthworm functions as well as the adverse effects that pollutants have had on earthworms as whole organisms.

## Methods

### Systematic literature search and screening

A systematic literature search was conducted in Web of Science on 18^th^ September 2025 with search terms related to earthworms, microbiota, the niche of the earthworm microbiota, and different pollutants (see Data S1 for the complete search terms). The process followed the PRISMA protocol (Preferred Reporting Items for Systematic reviews and Meta-Analyses)^31^.

In order to be suitable for the meta-analysis, articles needed to have 1) studied earthworms, 2) exposed the earthworms to a pollutant and 3) sequenced at least the microbiota of the earthworms after pollutant exposure. In the first step, the search results (n = 2,361) were manually screened for suitability based on their title and abstract using Abstrackr (abstrackr.com)^32^. We then screened the full texts of articles that passed this first stage of the screening (n = 276). Review articles were not included in the meta-analysis, but we screened the references cited in those reviews to identify additional suitable articles that were not already part of our literature search results. The whole screening process was performed by one screener (TMR). In total, full-text screening provided 159 suitable articles, including two papers identified from review articles. The screening workflow is summarized in the PRISMA diagram (Figure 1) with a more detailed workflow included in the supplementary (Figure S1).

**Figure 1.**
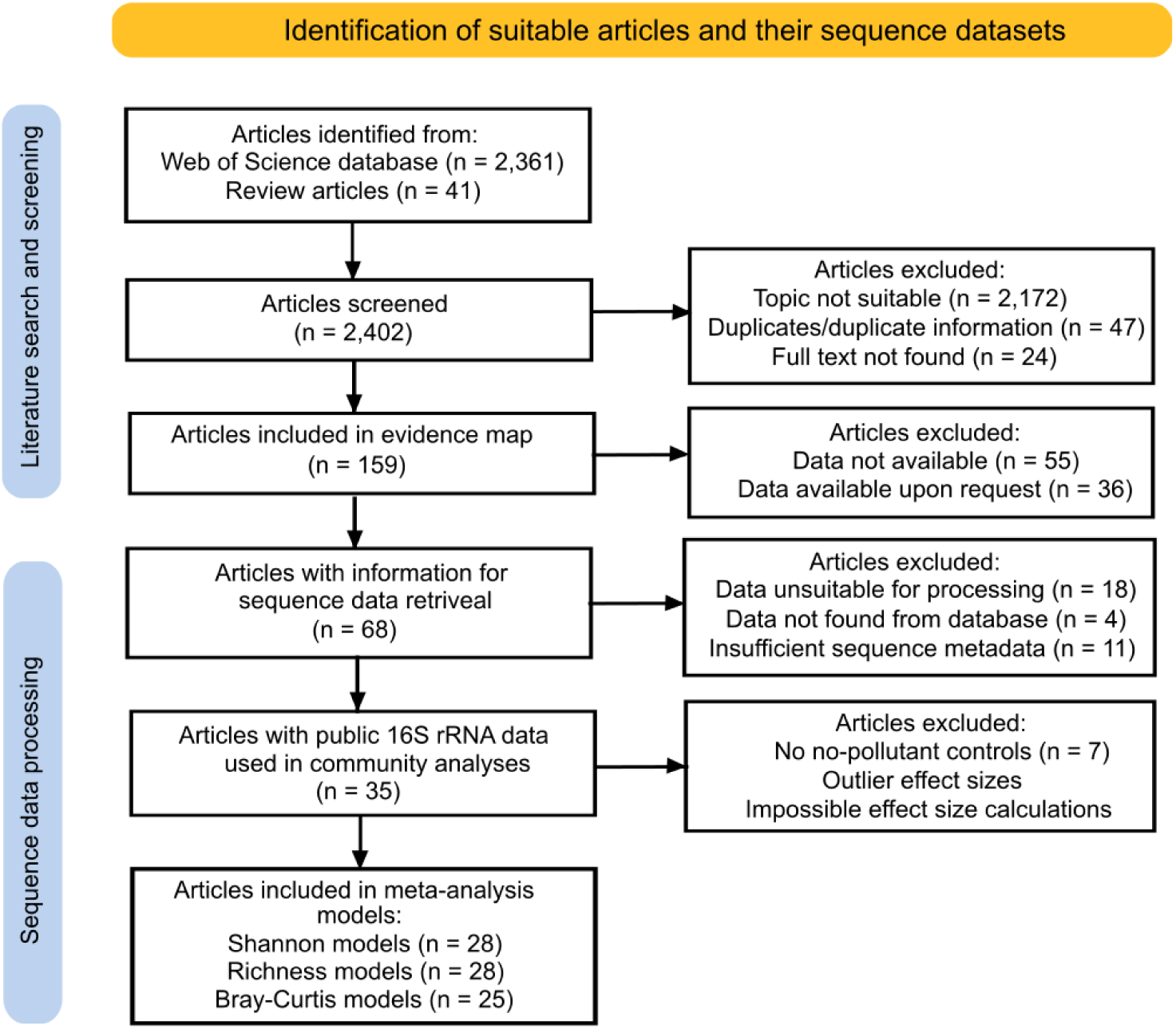
PRISMA 2020 diagram of the full workflow for article and sequence data identification. This diagram was created in Inkscape 1.4.2 (inkscape.org). A fully expanded PRISMA diagram is available in Figure S1.

### Metadata collection

Metadata was collected from 159 articles during full-text screening to create an evidence map. The main collected metadata and their definitions are summarized in Table 1.

**Table 1.** Metadata collected during full-text screening of each suitable article. Metadata used as predictors in the meta-analysis models are indicated with the predictor name inside the brackets.

| Meta-data variable (Predictor name) | Definition |
| --- | --- |
| Journal of publication | Journal the article has been published in. |
| Year of publication | Publication year of the article. |
| Pollutant group (Pollutant) | Which pollutant group(s) (Plastics, Fertilizers, Nanoparticles, Metals/half-metals, Organic wastes, Antibiotics, Pesticides, Organic wastes, and Other) did the tested pollutant(s) belong to. ‘Other’ included any pollutants that couldn’t be classified under any of the other groups. |
| Earthworm species (Species) | Scientific binomial of the earthworm species exposed to the pollutant(s) of interest during the experiments. |
| Earthworm origin (Origin) | Whether the earthworms were purchased from a breeder/grown in a laboratory, or whether they were collected from the wild. |
| Experimental location | Whether the experiments were carried out in laboratory or in the wild. |
| Microbiota source (Source) | Source of the microbiota (soil, gut, cast, gizzard, skin, whole earthworm etc.) |
| DNA extraction method | Which kit/method was used for DNA extraction |
| Sequencing platform | Which sequencing platform was used. |
| Amplification target and region | Which target gene(s) (16S rRNA, ITS, 18S rRNA) and its/their region(s) (e.g. V3-V4, V5) were amplified. The primers used for amplification were also recorded. |
| Diversity metrics | Which diversity (alpha, beta, gamma) had been calculated and did the calculated metric(s) measure taxonomic or phylogenetic diversity. |
| Data availability (Article ID) | Whether the article had made their sequence data public (Yes, no, available upon request). Any information for accessing sequence data (accession numbers or links) were recorded. Accession numbers were also used as IDs for the articles. |

### 16S rRNA metabarcoding data downloading

Of the 159 articles that were deemed suitable for full-text screening, 68 (42.8%) articles included enough information to retrieve their publicly shared sequence data. Out of these 68 articles, however, seven stored their data in non-standardized ways: three stored their sequence data outside of public databases (NCBI, ENA, CNCB, or DDBJ), and the data of four articles were stored in NCBI in a format we could not use for further analysis (e.g. sequence data stored as nucleotide data), and additionally, the data of one article was hosted on a sequencing platform which we were unable to access with our download tool. In contrast, we simply could not find the sequence data from 55 (34.6%) of the remaining articles, and, due to time constraints, at this stage, we did not contact the authors of the 36 (22.6%) articles that mentioned the data would be available upon request.

Due to the low amount of available metagenomic data (10 articles), ITS data (four articles), and 18S rRNA data (only one article), we decided to focus our analyses only on the available 16S rRNA metabarcoding data. When articles contained sequence data for multiple amplification targets, we only processed the 16S rRNA sequences. We collected the accession numbers from the remaining 50 articles (note, for this dataset, when focussing on 16S rRNA data, each article corresponds to a single BioProject in the NCBI database), however, only 46 articles had data that was downloadable. The full workflow of identifying suitable sequence data from the articles is summarized in the PRISMA diagrams (Figures 1&S1).

We used the Meta2Data pipeline found on GitHub (see https://github.com/LinyangSun/Meta2Data), to download and process the sequence data and their associated metadata using the high-performance computing (HPC) resources available from CSC – IT Center for Science, Finland (csc.fi). In brief, Meta2Data is a QIIME2^33^ based pipeline for automatic processing of public amplicon sequencing data. We first used the MetaDL function to search and download the metadata. We then extracted the raw sequence read data from the selected articles, and subsequently passed it into AmpliconPIP, which automatically detects and removes adapters and primers, conducts quality checks, and trims the sequence reads. The reads with sufficient quality scores were then denoised using DADA2^34^, whereas those with lower quality scores were processed using VSEARCH^35^. Finally, we assigned taxonomy to the sequence reads with the AmpliconTAXA function, using the Greengenes2 database^36^.

### Processing of public 16S rRNA sequence datasets

The pipeline downloaded and processed 16S rRNA sequence data from the 46 articles, which were then processed prior to analysis (Figures 1&S1). Unfortunately, 14 articles had submitted their sequence metadata insufficiently and we couldn’t distinguish unpolluted samples from polluted samples. We contacted the authors of these articles and received metadata from three. Therefore, sequence data from 11 articles were excluded. In addition, only samples taken from the earthworms or from their surrounding soil were included. Any vermicompost samples, organic waste samples or soil samples without earthworms were removed. Additionally, only freshly collected cast samples were retained, and any samples where microbes were specifically inoculated to the samples were removed. Typically, the research articles included multiple treatments, thus removal of such samples did not result in the full article being removed from the process.

The ASV-level data returned by the pipeline was agglomerated to genus level (from now on referred to as gASV to indicate genus-level ASV data). Any unannotated genera were labelled as unclassified with the lowest annotation level attached (e.g. Unclassified_Bacteria for those with only domain level annotation).

### Statistical analysis

We explored the overall effect of pollution of the earthworm microbiota and how the effects vary across different predictors through mixed effects meta-analysis models of the downloaded and processed 16S rRNA sequence data. A mixed effects modelling approach was used as samples within articles are not independent, and it is likely that sequences gained from a similar methodology are likely to be less heterogenous than across methodologies.

For the random effects component of the models, nested random effects were used. For random effect variables, sequencing platforms, amplification regions and primers were combined into a new variable, ‘sequencing’. Additionally, unique IDs were created for the articles and their ‘cases’ (discussed below). Accession numbers were used as article’s IDs and their cases were labelled numerically in increasing order.

Predictors (fixed effect components of the model) were identified from the collected metadata of the articles (Table 1). Some predictors were used as they were, but some were transformed for model fitting. The microbiota source was divided between external (soil, skin) and internal (gut, cast) levels for simplicity. One article tested co-exposure between two pollutant groups (i.e., metals/half-metals and plastics), and the effect of each pollutant could not be separated from the other, thus we placed these samples within the ‘Other’ pollutant group. As the microbiota of earthworms had been predominantly studied in *E. fetida*, the ‘*E. fetida* or other species’ predictor was created to only include two categories: the species *E. fetida* versus all other species.

In order to compare results across different articles, we chose to use the Hedge’s g effect size^37–39^, which is a unitless, standardized mean difference effect size. Hedge’s g was calculated using the metafor R-package^40^. Each article could provide multiple effect sizes, i.e, ‘cases’. For example, an article could test more than one pollutant, more than one earthworm species, or more than one microbiota source, where each would become its own case. Within each case, for samples in both the controls (i.e., unpolluted) and treatments (i.e., polluted) groups, the Shannon diversity, the gASV richness and the Bray-Curtis similarity index (1 – Bray-Curtis dissimilarity) were calculated using the ‘vegdist’ function of the vegan R-package^41^. Bray-Curtis similarity was calculated for Hellinger-transformed data, calculated with the ‘decostand’ function of the vegan package. For each of the three diversity metrics (Shannon, richness and Bray-Curtis), the mean, standard deviation, and sample size were calculated, and the Hedge’s g calculated. As effect size calculation requires a control and treatment, any articles without no-pollutant controls were excluded from the effect size calculation and therefore the meta-analysis. Additionally, articles or cases where effect size calculations were impossible or resulted in outliers were excluded (Hedges’ g > 5 or -5). The exclusion criteria for the meta-analysis models and the final number of articles and sample sizes in the meta-analysis models are summarized in the PRISMA diagrams (Figures 1&S1).

Using the metafor package, we used multi-level mixed effects models to analyse variations in the effects sizes of the three community metrics: Shannon diversity, richness, and Bray-Curtis similarity index. Random effects for all the models included the article ID nested within ‘sequencing’, and a case ID nested within article ID. For each model, the response variable was the Hedge’ g effect size from the Shannon, gASV richness or Bray-Curtis similarity. Within each of the three sets of models a null model was constructed that included no predictor variables, and in addition, univariate models were constructed using the following predictor variables: earthworm species, earthworm genus, *E. fetida* or other species, pollutant group, DNA extraction method, earthworm origin, and microbiota source. Within each set of models, each univariate model was compared to the null model, using a Wald-Type test (‘anova’ functions in metafor). The predictor variable was not considered significant if p > 0.05. Orchard plots were created for significant predictors with the orchard_plot function of the orchaRd R-package^42^. To check for publication bias, funnel plots of the entire dataset were created using the ‘funnel’ function in the metafor package.

To further explore possible differences in the gut microbiota (i.e., internal microbiota) associated with *E. fetida* versus all other species, we used the 16S rRNA data at the class level to identify the most abundant microbial taxa in these two groups of species. Identifications were made for controls and treated samples separately to test whether the taxa changed under pollutant exposure. Unlike for the meta-analysis models, the articles without no-pollutant controls were included in these calculations. The final number of articles and sample sizes are summarized in the PRISMA diagrams (Figure 1&S1).

All statistical analyses were performed and figures created in R 4.5.2^43^. To visualize the metadata collected from the articles and to create meta-analysis models from processed sequence data, the R-package tidyverse^44^ and associated packages were used. Some figures were further modified with Inkscape 1.4.2 (inkscape.org). The output from the Meta2Data pipeline was read into R with the qiime2R package^45^.

### Data availability

All code and data (including meta-data and effect sizes) used in this study are publicly available in Zenodo (https://doi.org/10.5281/zenodo.22670569).

## Results

### Evidence map

We created an evidence map based on the metadata from 159 articles testing the effects of pollutants on microbial communities associated with earthworms (Figures 2,3&4). The earliest publication dates from 2011, but the field has experienced a very clear increase in publications since 2018 (Figure 2A), with an obvious increased interest for this field in China since this date (n = 122, Figure 2B). Most of the articles are published in the Journal of Hazardous Materials (n = 33), Science of The Total Environment (n = 21), Environmental Pollution (n = 18) and Chemosphere (n = 11) (Figure S2).

**Figure 2.**
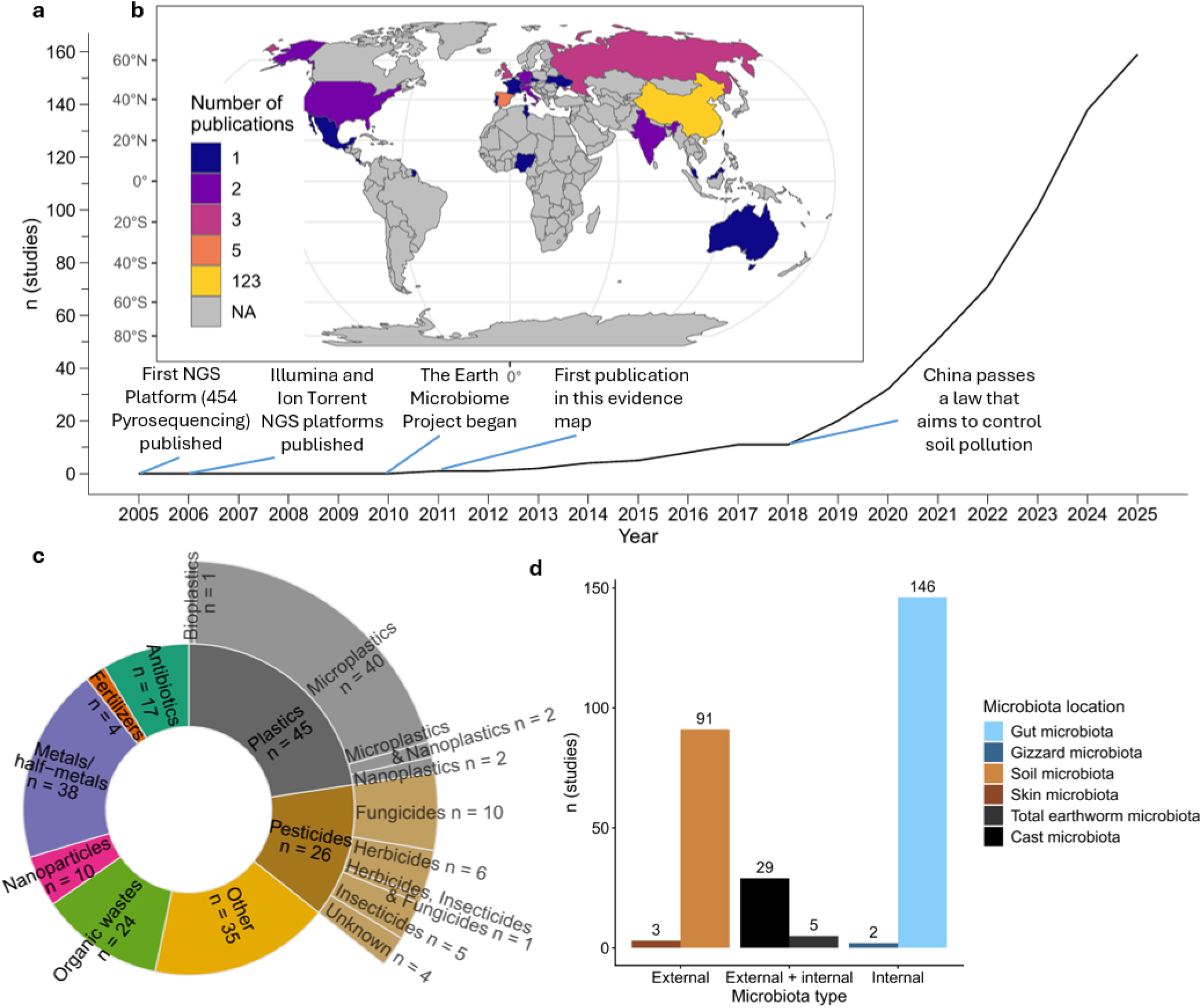
A) Cumulative number of articles over time and B) geographical locations of the articles. Geographical locations of the papers were deduced from the main institutions of the authors and methodology of the experiments. The C) pollutant group(s) and D) microbiota sources used in the articles. Total earthworm microbiota refers to the microbiota found in the whole earthworm, in most cases taken from crushed earthworms. In all figures, n corresponds to the number of articles. Articles sampling the microbiota from multiple sources and testing multiple pollutant groups will be duplicated across figures C and D.

The most commonly tested pollutant group were plastics (45 articles) (Figure 2C), with most articles investigating the effect of microplastics (40 articles). Metals/half-metals were the second most studied pollutant group (38 articles). And perhaps surprisingly, only four articles tested the effect of soil fertilizers.

Almost all articles (n = 146) focused on the effects of these pollutants on the earthworm gut microbiota (Figure 2D), and many articles (n = 91) sampled the soil microbiota in addition to the earthworm microbiota. The microbiota of cast was also studied in 29 articles. Only 10 articles sampled earthworm microbiota from other sources than the gut or casts, including three articles that sampled the skin and two articles that focused on the gizzard (an organ responsible for grinding ingested soil^46^). No articles sampled the microbiota of nephridia or cocoons.

In most of the articles (n = 102) the effects of pollutants were tested on the earthworm species *Eisenia fetida* (Figure 3), but there were 24 species represented in the dataset, with a further two articles not identifying the earthworm species, and one article only identifying the earthworm genus. Regardless of the species, most of the earthworms used in the experiments were purchased from breeders, or were lab-grown (117 articles), and a minority of articles focused on wild specimens (23 articles). Similarly, almost all the articles (n = 150) were conducted in laboratory or artificial environments and only a handful (n = 9) were field experiments. Unfortunately, an additional 20 articles didn’t mention the origin of their study organisms.

**Figure 3.**
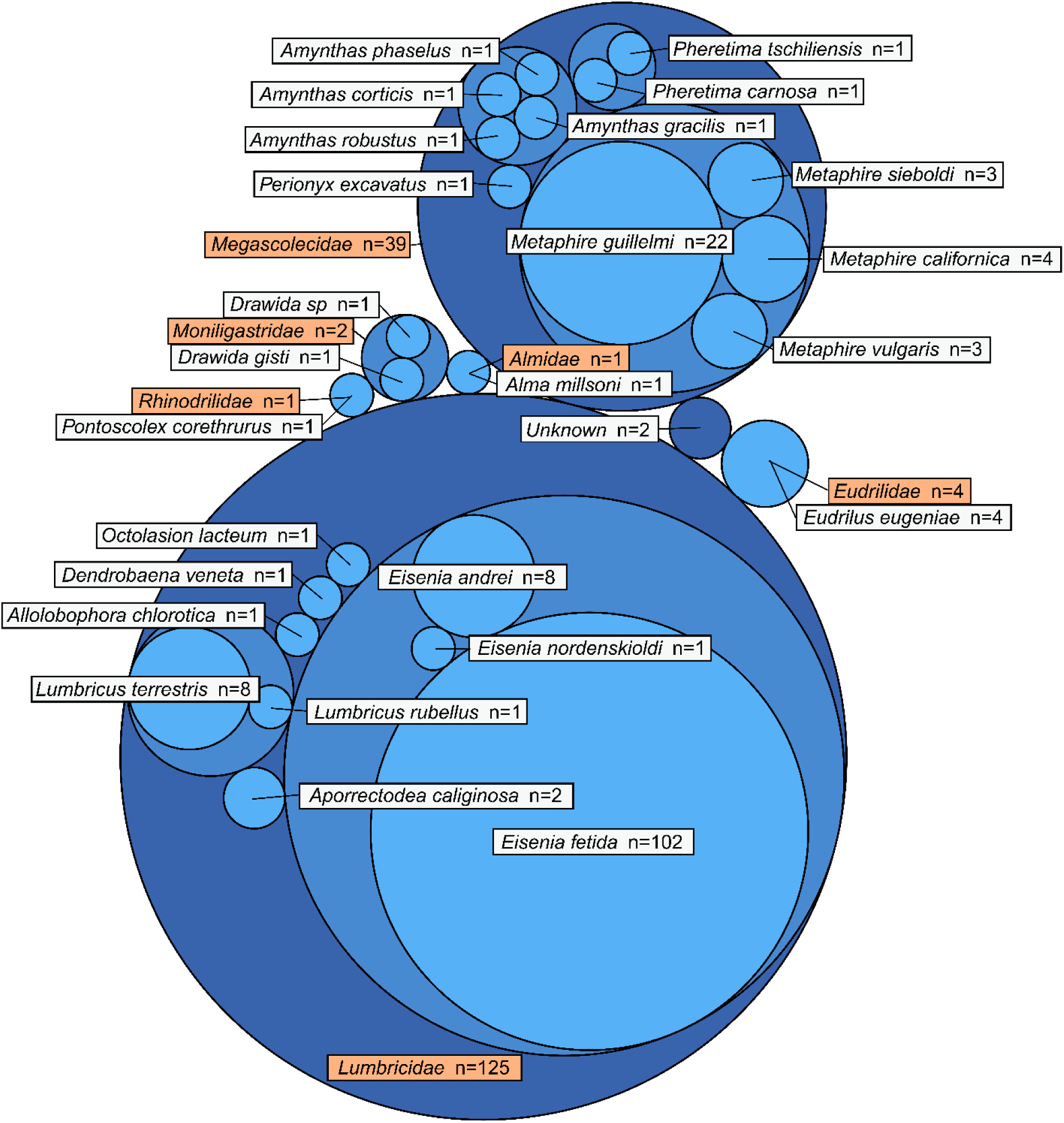
A hierarchical circle packing chart of earthworm taxa with hierarchy proceeding from family level to genus level to species level. Species names labelled with white backgrounds and family names with orange backgrounds. Genus labels not shown. Circle size and n correspond to the number of articles. Articles that experimented with multiple species are duplicated across the figure.

The most commonly used approach to characterize earthworm microbiota was through 16S rRNA metabarcode sequencing (146 articles) (Figure 4). The 16S rRNA sequencing allows the characterization of bacterial and archaeal communities only. The V3-V4 hypervariable regions of the 16S gene was the most common target (70 articles). In contrast, 10 articles targeted the *ITS* gene to characterize fungal communities, and only two articles targeted the 18S rRNA gene to characterize protist communities. Finally, 12 articles used shotgun metagenomics to target all genetic material from the microbial communities. Using the sequence data, most articles calculated metrics of taxonomic alpha and beta diversity of microbial communities, with minimal metrics of phylogenetic diversity (Figure S3).

**Figure 4.**
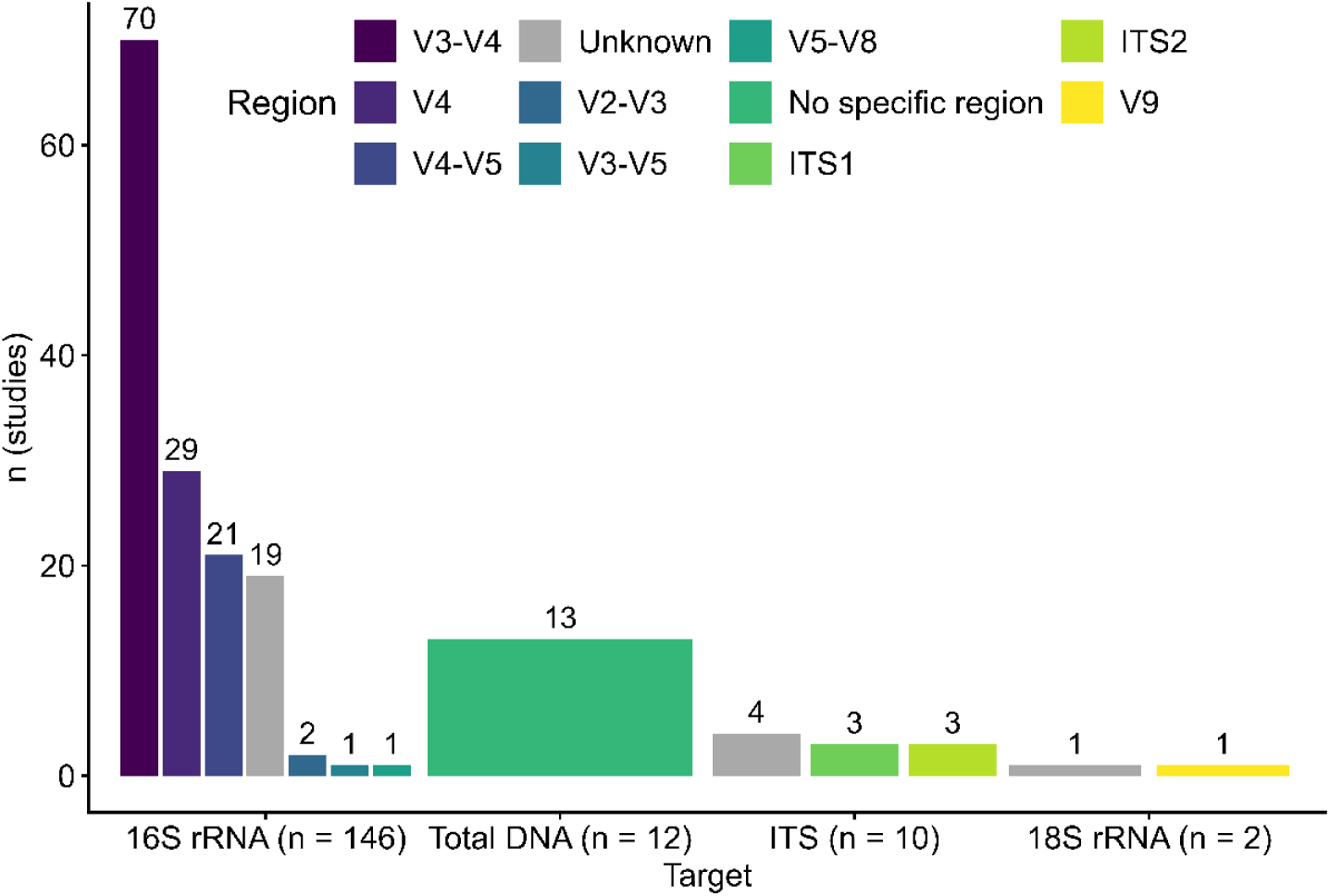
Amplification targets (16S rRNA, total DNA, ITS and 18S rRNA) and their specific regions sequenced in the articles, where n corresponds to the number of articles. Bar width does not convey information. Articles sequencing multiple targets or regions are duplicated across the figure.

### Meta-analyses of earthworm microbiota responses to pollution

Based on the Null models, there was no overall significant effect of pollution on the Shannon diversity (estimate = 0.0574, S.E. = 0.1501, p-value = 0.7023, z-value = 0.3822) and gASV richness (estimate = 0.1822, S.E. = 0.1434, p-value = 0.2040, z-value = 1.2702) of the earthworm microbiota. Pollution however significantly decreased the Bray-Curtis similarity index of the earthworm bacterial communities (estimate = -0.4604, S.E. = 0.1616, p-value = 0.0044, z-value = -2.8485). There was no strong evidence of publication bias within the compiled data (Figure S4).

In contrast, variations in effect sizes were significantly explained by several predictors. The predictors ‘type of pollutant’ and ‘source of the microbiota sample’ both significantly explained additional variation in the Shannon diversity models. These two predictors did not however affect the gASV richness and Bray-Curtis similarity models (Table 2). The predictor ‘DNA extraction method’ also significantly explained variation in all three models, however, as all factor levels in this predictor were data-poor (Figure S5), these effects were not further investigated as the model lacked robustness.

**Table 2.** Univariate model comparisons with the null model using Hedge’s g calculated from Shannon diversities, gASV richness and Bray-Curtis similarity as the response variable. Within the univariate models, predictors were the pollutant type, earthworm species, earthworm genus, earthworm origin, microbiota source (e.g. which tissue), or whether the species was E. fetida or another. The effects of the bolded predictors were significant with p-value < 0.05.

| Response variable | Predictor | Degrees of freedom | p-value | AIC | QE | I<sup>2</sup> |
| --- | --- | --- | --- | --- | --- | --- |
| Shannon | Null | 4 | NA | 237.4340 | 203.4131 | 68.23915 |
|  | **Pollutant** | **11** | **0.0207** | **234.9097** | **160.6970** | **62.16935** |
|  | Species | 15 | 0.6296 | 250.5173 | 185.5902 | 71.56179 |
|  | Genus | 13 | 0.4772 | 246.8571 | 187.8922 | 70.18786 |
|  | *E. fetida*/other | 6 | 0.3758 | 239.4765 | 202.7248 | 69.13807 |
|  | Origin | 6 | 0.5574 | 240.2651 | 202.9705 | 68.44205 |
|  | **Source** | **5** | **0.0222** | **234.2058** | **195.2241** | **66.61846** |
| gASV richness | Null | 4 | NA | 210.8838 | 157.2146 | 58.46418 |
|  | Pollutant | 11 | 0.0820 | 212.2680 | 128.7385 | 53.59085 |
|  | Species | 15 | 0.6043 | 223.6927 | 142.9070 | 62.78748 |
|  | Genus | 13 | 0.4618 | 220.1453 | 144.4629 | 61.26969 |
|  | *E. fetida*/other | 6 | 0.1720 | 211.3636 | 153.0196 | 57.79123 |
|  | Origin | 6 | 0.0734 | 209.6594 | 153.4034 | 56.3435 |
|  | Source | 5 | 0.4999 | 212.4288 | 156.7650 | 58.65763 |
| Bray-Curtis | **Null** | **4** | **NA** | **194.2181** | **141.7475** | **56.0495** |
|  | Pollutant | 11 | 0.0746 | 195.3197 | 113.8450 | 53.97918 |
|  | Species | 15 | 0.1582 | 200.6571 | 103.2568 | 52.81501 |
|  | Genus | 13 | 0.1119 | 197.9145 | 105.4010 | 50.63566 |
|  | *E. fetida*/other | 6 | 0.1104 | 193.8113 | 133.4221 | 54.93325 |
|  | Origin | 6 | 0.5713 | 197.0984 | 137.8517 | 57.80713 |
|  | Source | 5 | 0.6338 | 195.9912 | 141.6648 | 57.22395 |

The predictor ‘pollutant type’ also had a significant effect on Shannon diversity, with specimens treated with antibiotics showing a significantly increased Shannon diversity (estimate: 2.7027, S.E. = 0.7826, p-value = 0.0006, z-value = 3.4537) (Figure 5A). In the model using the predictor ‘microbiota source’, neither of the factor levels were significantly different from zero (Figure 5B), but they were significantly different from each other, with the Shannon diversity associated with the earthworm’s internal microbiota being significantly higher (estimate: 0.2378, S.E. = 0.1672, p-value = 0.1549, z-value = 1.4224) than that of the external microbiota (estimate: -0.4320, S.E. = 0.2417, p-value = 0.0739, z-value = -1.7873).

**Figure 5.**
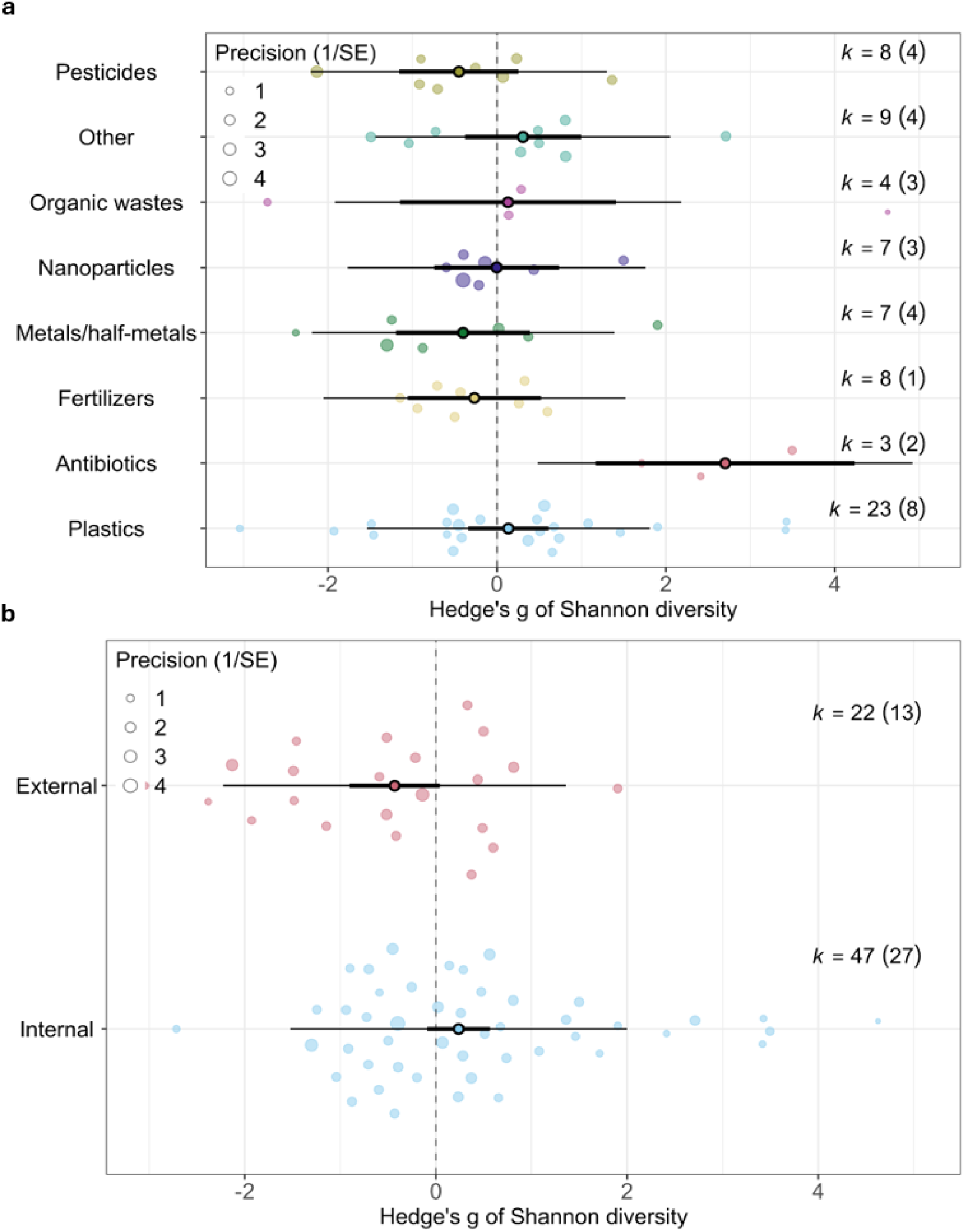
Orchard plots for the univariate meta-analysis models. A) Pollutant model and B) microbiota source model of Shannon diversity. Central dot indicates the overall effect of the factor level of the predictor on the effect size. Dark bars indicate the 95% confidence interval, and thin bars indicate the S5% predictive interval. Predictor factor levels are considered significantly different from zero when the confidence interval does not overlap with zero. K indicates the number of cases and the number of articles is given in parentheses.

### Most abundant bacterial taxa for *E. fetida* versus all other earthworm species

We identified and compared the most abundant microbial taxa associated with internal tissues of *E. fetida* versus all other earthworm species, between control and treated groups. The most abundant bacteria colonizing *E. fetida* were from the class *Gammaproteobacteria,* while class *Bacilli* was the most abundant in the all-other species-group. Pollutant exposure did not affect the identity of the most abundant bacterial taxa in either *E. fetida* or in the all-other species-group. The list of the 20 most abundant taxa found in association with all samples are given in figures S6&7, while the corresponding read counts can be found in figures S8&9. The samples represented a good coverage of different genera (Figure S10).

## Discussion

We extracted and analysed the metadata from 159 articles investigating the effects of pollutants on the microbiota of earthworms. We complemented this systematic review, with a meta-analysis of the 16S rRNA metabarcoding sequence data publicly available from only a subset of these articles. Unfortunately, there was a distinct lack of articles with available sequence data, and clear biases within the wider literature. The majority of the literature used *E. fetida* as a model organism and focused on the effects of plastics on the gut microbiota, and thus, the current literature may not be able to provide an accurate representation of the overall effects of pollutants. Consequently, because effect sizes were highly variable, we have no convincing evidence that pollutants have a significant effect on the Shannon diversity or richness index of the earthworm microbiota. One specific pollutant group, ‘antibiotics’, showed significantly increased Shannon diversity, but we were unable to account for variation in effect sizes between the treatment groups. We also found Shannon diversity to be significantly different between internal and external samples, although neither were significantly different from zero. Pollutant exposure resulted in the overall decrease in Bray-Curtis similarity indicating changes in the community compositions after exposure, with microbiota samples being more dissimilar to each other than in non-polluted samples, but we found no moderating effects to explain this.

### Impact of pollution of earthworm microbiota

Across the full body of literature, plastics were the most studied pollutant (45 articles) and, mirroring the wider microbiota research^47^, most plastics tested were microplastics (40 out of 45 articles). However, despite plastics being a widely distributed pollutant^47^, we found no indication that they have a significant effect on the earthworm microbiota. Although plastics can accumulate over time within many organisms^48^, this does not appear to be the case for earthworms^49^, which could explain the lack of significant effect as the microbiota is able to recover after the exposure. While the median number of exposure days of the studies included (median days = 20) is in line with OECD guidelines for earthworm ecotoxicity tests (which recommend 7 and 14-day incubation times^50^, or 28-day incubation times for *E. andrei* and *E. fetida* specifically^51^, we are unable to determine whether the dose of plastic pollution was large enough to impact the effect of such pollution, or any pollution type, on the earthworm microbiota. In contrast, we recommend additional studies on underrepresented pollutants, such as fertilizers. As the amount and rate of fertilizer application has increased substantially in the past decades^52^, and has had significant impact on invertebrate communities^19^, especially at the highest levels^53^, understanding the impact of fertilizers on earthworm microbiota is important^54^.

Most of the articles (n = 122) have been conducted in China, where the agricultural sector is extremely large^55^, and industrialization is still growing^56^. Agriculture is the largest contributor to global metal emissions out of all anthropogenic sources, due to the application of agrochemicals^57^, while industrialization contributes significantly to metal pollution of soils^57^. Given the generalized soil degradation that the country faces^55^ and the importance of maintaining soil health for the provision of food^58,59^, China has recently, in 2018, passed ‘The Law of the People’s Republic of China on Prevention and Control of Soil Contamination’. This new law enforces the protection of the soil environment and aims to control contamination (https://english.mee.gov.cn/Resources/laws/environmental_laws/202011/t20201113_807786.shtml; Article 1). The impact of this law reflects in the priorities of the scientific research conducted in this country, with the number of earthworm microbiota studies in China within the 5 years following 2018 rising from 4 to 81, which also explains the strong exploration of the impact of metal pollution (n = 38) on earthworm microbiota within the literature. While there is evidence that metals can be highly detrimental to earthworms in terms of their richness and diversity^60,61^, we found no significant effect of metal exposure on the earthworm microbiota, highlighting the relative resiliency or adaptation of the microbiota to metals detected in previous studies^62–64^. Adequately testing the specific impacts of metal pollution on, for example, different species or different organs^65,66^, would require additional data covering a wider variety of species, or organ tissues, as treatment groups.

Antibiotics often decrease microbial diversity within organisms, as shown in humans^67^ and invertebrates^68^. In earthworms, previous studies have shown that the microbiota typically contains lower antibiotic-resistant gene-carrying microbes^69^, making the earthworms’ microbiota possibly more prone to loss of microbial species or microbial abundance following antibiotic application. Against our expectation, we recorded that antibiotic treatments were positively affecting microbial diversity in earthworms. Although this result was only supported by two articles with limited data in our meta-analysis, it is possible that the antibiotics might have suppressed some dominant microbial taxa while allowing for new opportunistic taxa (from the soil) to be established or increase in abundance^70^, especially if the newly established taxa have developed antibiotic resistances under antibiotic pressure^71^. Nonetheless, as of now, we do not consider this result to be robust enough to confirm any true biologically meaningful conclusion.

### Taxonomic representation

To date, the meta-analysis of global patterns in earthworm’s microbiota, could only be performed using bacterial 16S rRNA data – as ITS and 18S rRNA amplicon sequencing is only available from four and one unique article, respectively. The abundance of bacteria is higher in the earthworm gut^72^, and thus the predominant use of the 16S rRNA gene to identify prokaryotes is to be expected^73^, but as both fungi and protists are a part of the earthworm diet due to their presence within the ingested soil^11,74^ we hope that future studies will consider multiple amplification targets.

The 24 species described in the literature included in this meta-analysis only represent a very small fraction of the known earthworm species biodiversity. For example, the *Megascolecidae,* which is the largest family with 2347 species^75^, was the most represented family in our analyses, with only 11 species. Similarly, only nine of the 688 species described within the *Lumbricidae* family, including *E. fetida*, were represented in the present study. While the *Acanthodrilidae* family, which is the second largest earthworm family, wasn’t represented at all.

Historically, the earthworm species *Eisenia fetida,* has been described as highly sensitive to pollutants^76,77^, and has thus been used in numerous ecotoxicity studies. Out of the 24 earthworm species used in our meta-analysis, *E. fetida* was the most common target (102 articles). In contrast with historical expectations, our results, and those of other recent studies^78–80^, indicate that this species may not as sensitive to pollutants as previously thought. A meta-analysis comprising 44 ecotoxicity experiments found that *E. fetida* is in fact one of the least sensitive species to pesticides^81^. Similarly, two studies comparing *E. fetida* to *Metaphire guillelmi’*s_responses to pollutants (di(2-ethylhexyl) phthalate (DEHP) and polycyclic aromatic hydrocarbon (PAH) pyrene) found smaller decreases in gut microbial diversity in *E. fetida*^80,82^.

From the downloaded 16S rRNA amplicon data used in our study, *Gammaproteobacteria* was the most abundant bacterial class for *E. fetida* in both treated and control groups, but was not the most abundant class in other species. *Gammaproteobacteria* have been shown to increase in abundance after exposure to the heavy-metal chromium^83^ and PAHs^84^, potentially resulting in species specific responses to pollution. Given the large percentage of unclassified microbial taxa within our study, at both class and family level (Figures S6&7 respectively), more studies on the microbial compositions of different earthworm species are needed to better understand these differences in sensitivity. But as *E. fetida*, and perhaps it’s microbiota, may be somewhat resilient to pollution, using it as a model species in pollutant studies may not give an accurate representation of how pollution impacts more sensitive species.

Noticeably, most of the earthworms used in the studies (n = 117) were commercially bought or reared in the laboratory, and only a few articles (n = 23) documented data and results from wild specimens. Because of their simplified environment, captive specimens usually carry a lower microbial alpha diversity than individuals from the wild. This observation was illustrated in mice^85^, cockroaches^86^ and *Drosophila* flies^87^. More pollution studies should be conducted using wild earthworms and a variety of species considering that diversity differences exist between species and possibly between laboratory-grown and wild individuals, which could greatly influence the results found.

### Microbiota tropism

Unsurprisingly, the most extensively studied microbiota in earthworms (146 out of 151 articles) was that of their gut, which plays an important role in organic matter degradation, fatty acid production and detoxification^9^. In contrast, the earthworm skin microbiota and gizzard were sampled and characterized in only three and two articles, respectively, despite their importance in earthworm biology. The earthworm skin microbiota plays an important role in earthworm immunity^9^ while the gizzard is a soil grinding organ^88^, and thus a major part of the earthworm digestive system.

There were no articles that had sampled the cocoon or nephridia microbiota. Earthworm cocoons mostly inherit their microbes vertically from their parents during cocoon formation^89^, with a small amount originating horizontally from the soil^13^. The nephridia microbiota is one of the most important inherited components as it consists of conserved symbionts important in the host’s nitrogen recycling, immunity and detoxification^9^. These nephridia symbionts and earthworms show strong evidence for millions of years of host-symbiont co-evolution^90,91^ that may unfortunately be disrupted by pollution, as pesticides such as imidacloprid, benomyl and metribuzin have been shown to decrease the relative abundance of *Verminephrobacter* in the gut, one of the main symbionts of the nephridia^92^.

Overall, compared to the gut microbiota^93^ there is a distinct lack of knowledge of other important organs in the earthworm microbiota. We recommend more studies to focus on the skin and gizzard microbiota to determine how pollution may be affecting earthworm immunity and digestion before the gut^88^, as well as understanding the link between the microbiota of parents and their offspring through the nephridia and cocoons.

### Temporal trends

The increase in earthworm microbiota studies over times aligns with the development of technology and the wider research field (Figure 1A). The first earthworm microbiota study in this evidence map was published in 2011, shortly after the development of the more cost-effective next-generation sequencing (NGS) platforms, Illumina and Ion Torrent in 2006 and 2010 respectively^94^. The cost-effectiveness of the method is also reflected in which region of the 16S rRNA gene was sequenced - the V3-V4 region was the most common, sequenced in 70 articles, the length of which corresponds to some of the most cost-effective sequencing platforms^95^. Additionally, the launch of the Earth Microbiome Project (EMP) in 2010, which aimed to explore the microbial diversity from hundreds of thousands of samples around the globe^96^, highlights the growing interest in microbiota studies around that time, most likely inspired by the development of sequencing techniques. As the number of publications started to rapidly increase only after 2018, the increase is probably driven by the passing of the ‘The Law of the People’s Republic of China on Prevention and Control of Soil Contamination’ in China, which likely increased funding for research projects studying pollutant effects on the soil fauna, including earthworms.

## Data Availability

Unfortunately, while many journals require data to be open-access^97,98^, we observed that authors do not always take the time to transform their data into a standardized format, and mis-labelling becomes common, thus hindering data reusability^99^. Consequently, sequence data was not available for 55 articles in our metadata literature, while an additional 14 studies included samples incorrectly or insufficiently labelled, and 36 stated that the data would be only available upon request. These practices increase the risk of data loss in the long-term^100^ and go against good data-sharing policies from most journals and funders^101,102^. Only 68 of the 159 articles had made their data public. We join with others in promoting open-access data using FAIR principles^103^, to make research more reproduceable and optimize the use and re-use of one’s dataset, thus preventing valuable data from being lost and accelerating scientific discovery.

## Conclusions

Despite a recent increase in the number of publications, the field studying pollution effects on the earthworm microbiota is still young and hindered by a lack of publicly available sequence data and biases in the study designs. As while we didn’t find strong effects of pollution on the earthworm microbiota, we expect microbiota responses to be host specific as different earthworm species have been found to harbour a different composition of microbes. Consequently, we suspect that if we were able to include a higher diversity of species there could potentially be a shift in the non-significant results seen in this study. Current studies focus mainly on the digestive system, but as the earthworm nephridia, cocoons and skin are important in reproduction and immunity, their microbiota could help better understand findings from traditional ecotoxicology studies exploring earthworm survival and reproduction. Unfortunately, although NGS-sequencing has been around for two decades now, there remains a lot we don’t know about earthworm microbiota, reflected in the many undescribed taxa present in our dataset. Therefore, we propose more studies to characterize the microbial species in earthworms to lay a better groundwork for understanding changes even at lower taxonomic levels. We hope that by identifying the biases and gaps in the field, our results will help guide future studies towards more variety and open data principles, as this data provides valuable insights into the ecology of earthworms and their associated microbes under changing environmental conditions.

## Author contributions

TMR, AD and HRPP conceived the idea, TMR collated all data, LS provided software for accessing and downloading sequence data, TMR analysed all data and produced all figures. The initial draft of the manuscript was written by TMR and HRPP, and all authors provided feedback on subsequent drafts.

## Acknowledgements

We thank CSC – IT Center for Science, Finland for their generous resources for downloading and processing the sequence data. We also thank the authors who kindly provided their sequence metadata for the analyses.

