## Supplementary Material for "A meta-analysis on the effects of pollutants on the earthworm microbiota"

Data S1. Search word string used in the Web of Science database on 18^th^ September 2025

earthworm* AND (microb* OR "gut flora" OR "intestinal bacteria" OR "bacterial communit*" OR holobiont* OR *symbiont* OR commensal* OR fung* OR bacteria* OR prokaryote* OR micro$organism*) AND (gut OR intestin* OR “intestinal tract” OR “digestive tract” OR “gut content*” OR “gut soil” OR skin OR epiderm* OR nephridia* OR cuticle OR gizzard OR mouth OR esophagus OR pharynx OR feces OR cast OR manure OR earthworm$associated OR host$associated OR coelom* OR cocoon* OR clitellum) AND (pollut* OR contamin* OR toxi* OR metal* OR chemical* OR asbestos OR radionuclide* OR radioactiv* OR pharmaceutic* OR “emerging contamin*” OR “synthetic organic chem*” OR “personal care product*” OR *plastic* OR “polycyclic aromatic hydrocarbon*” OR pesticide* OR herbicide* OR fungicide* OR molluscide* Or nematicide* OR insecticide* OR *chemical* OR “oil spill*” OR “brine spill*” OR “petrol spill*” OR mining OR smelting OR “industrial activit*” OR “waste disposal” OR wastewater OR sludge OR sewage OR antibiotic* OR “polychlorinated biphenyl*” OR nano* OR “nitrogen deposition” OR “nutrient deposition” OR “atmospheric deposition” OR *eutroph* OR fertili* OR “nutrient* enrichment” OR “nutrient pollut*” OR detergent* OR biocide* OR “organic pollutant*”)

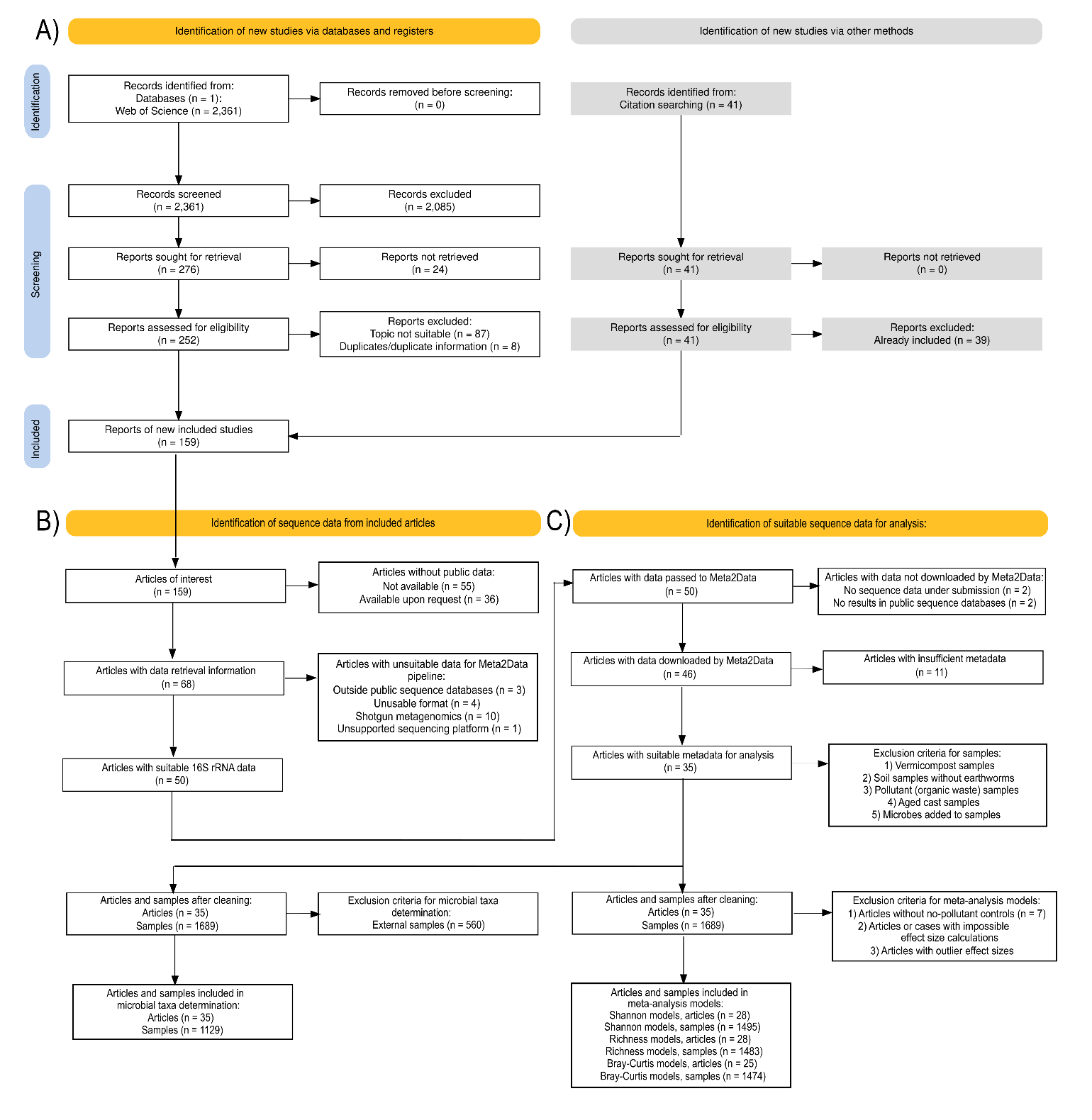
*Figure S1. Full extended PRISMA 2020 diagram of the workflow for A) literature screening*^1^*. This diagram was expanded further in Inkscape 1.4.2 (inkscape.org) to contain the workflows for B) sequence data identification and C) cleaning of the identified and processed sequence data.*
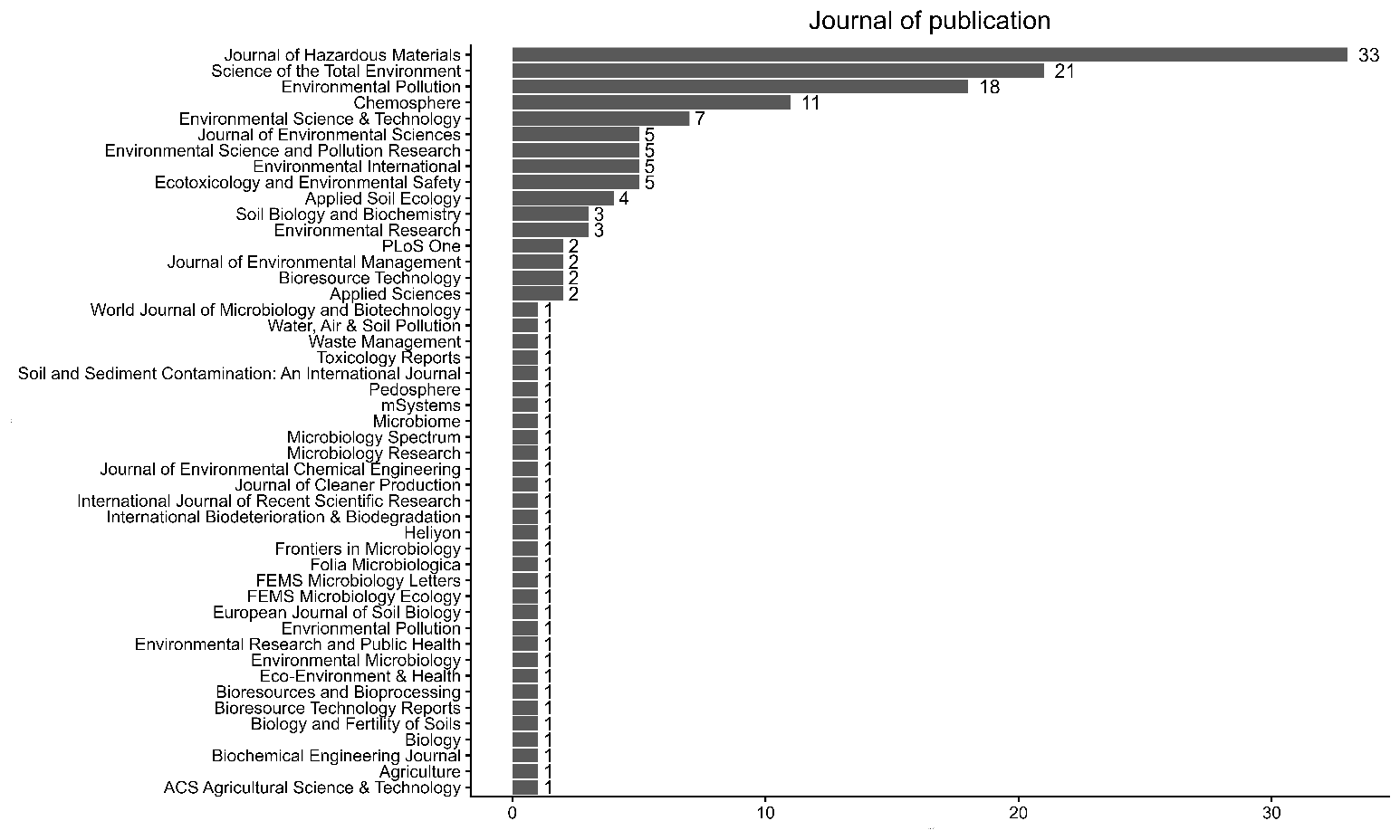

*Figure S2. Publication journals of the articles where n corresponds to the number of articles.*

*Figure S3. Sankey plot for diversity metrics calculated in the articles where n corresponds to the number of articles.*

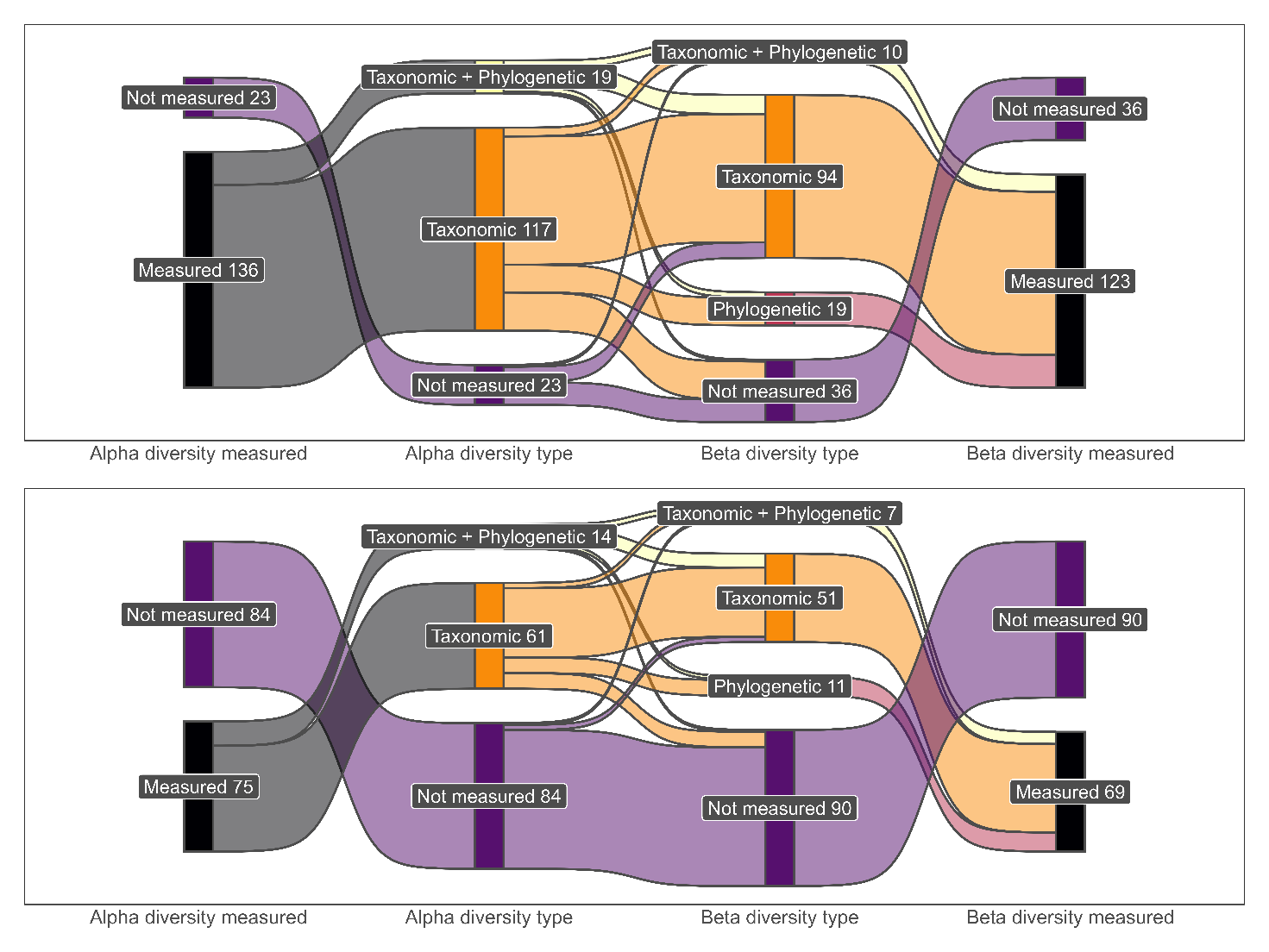

A

B

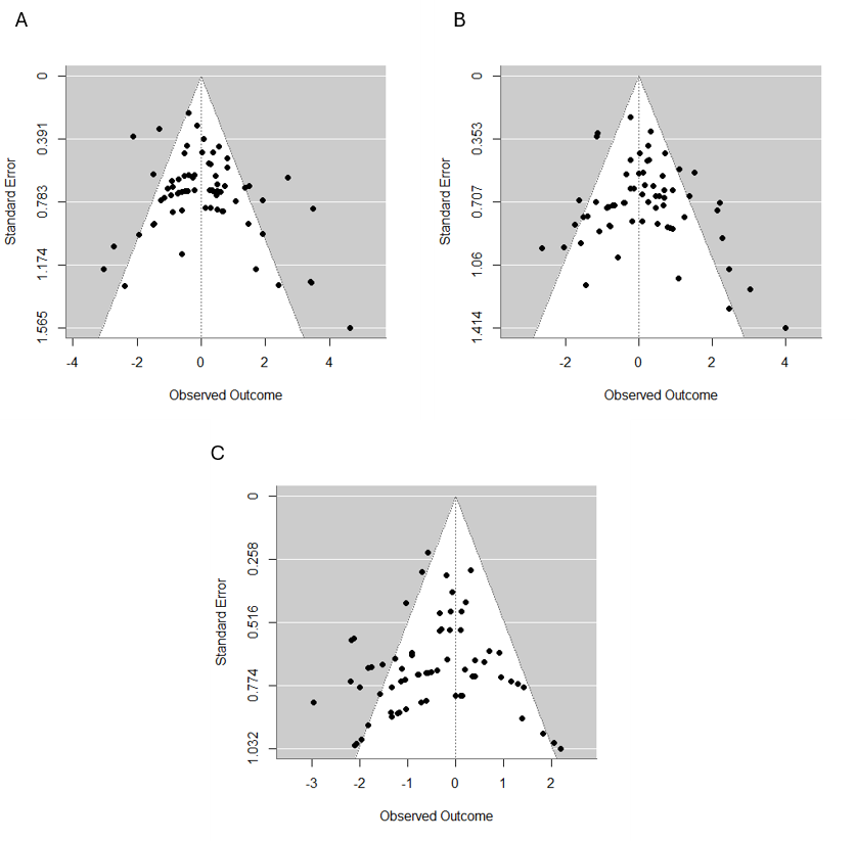
*Figure S4. Funnel plots of the A) Shannon diversity model, B) richness model and C) Bray-Curtis similarity model for detecting publication bias of the articles.*

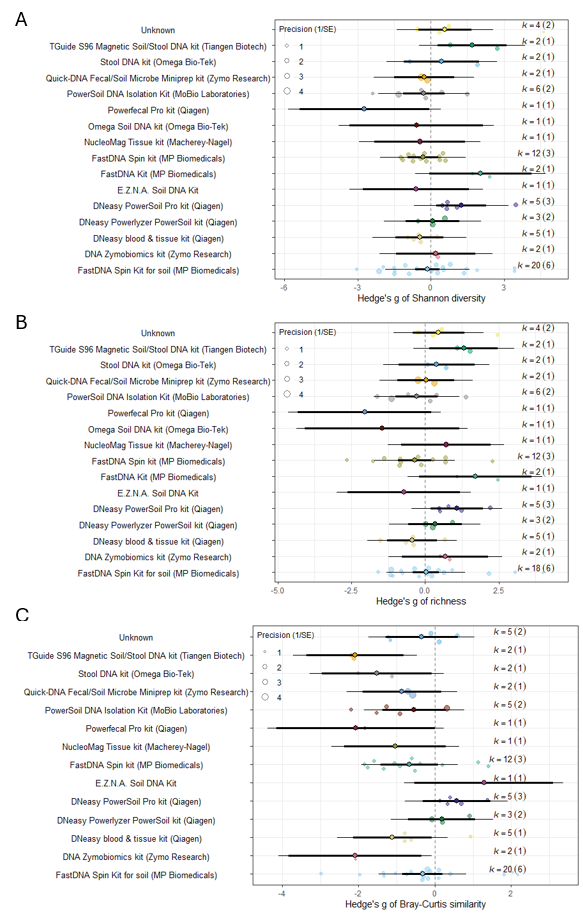
*Figure S5. Orchard plots of the DNA extraction univariate models for the A) Shannon diversity, B) richness and C) Bray-Curtis similarity meta-analysis models. Central dot indicates the overall effect of the predictor on the effect size. Dark bars indicate the 95% confidence interval, and thin bars indicate the predictive interval. Predictor levels are considered significantly different from zero when the confidence interval does not overlap with zero. K indicates the number of cases and the number of articles is given in parentheses.*

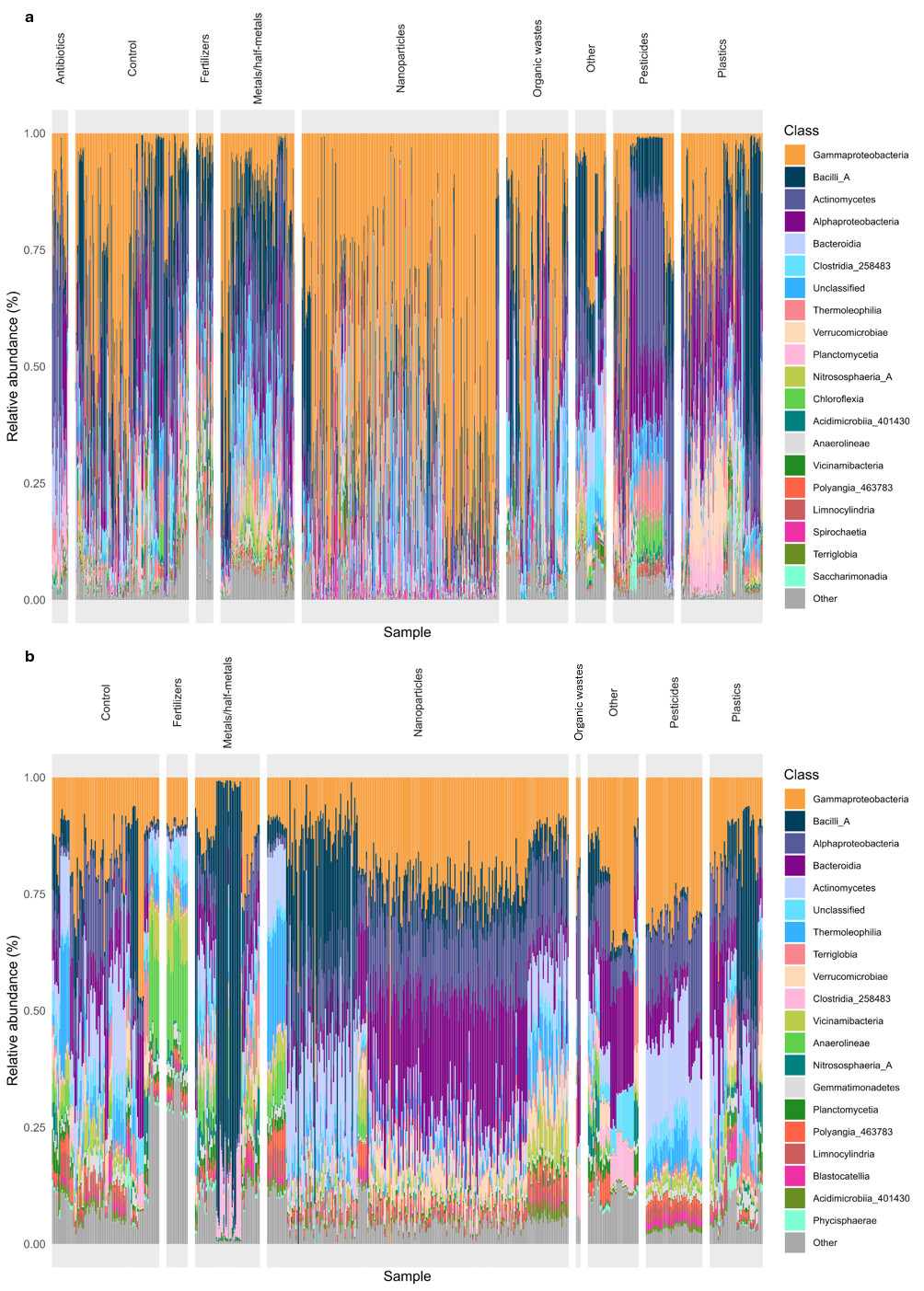
*Figure S6. Relative abundances of the top 20 microbial taxa for the different pollutant groups A) in the internal (gut, cast) samples and B) in the external (skin, soil) samples at class level. Remaining taxa grouped together as ‘Other’. All unannotated taxa at the class level were grouped together as ’Unclassified’ to visualize the dominance of unclassified taxa in the samples.*

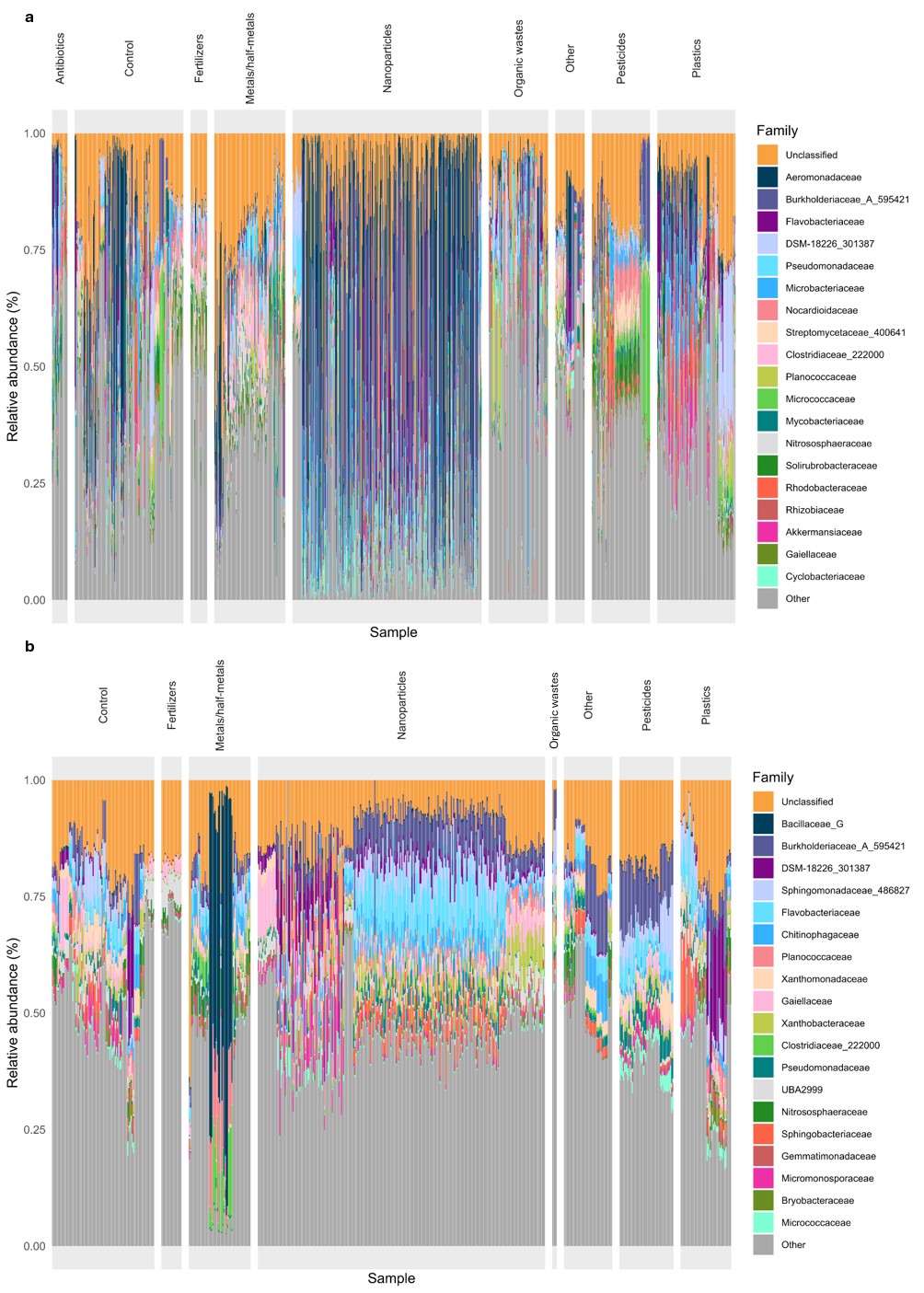
*Figure S7. Relative abundances of the top 20 microbial taxa for the different pollutant groups A) in the internal (gut, cast) samples and B) in the external (skin, soil) samples at family level. Remaining taxa grouped together as ‘Other’. All unannotated taxa at the family level were grouped together as ’Unclassified’ to visualize the dominance of unclassified taxa in the samples.*

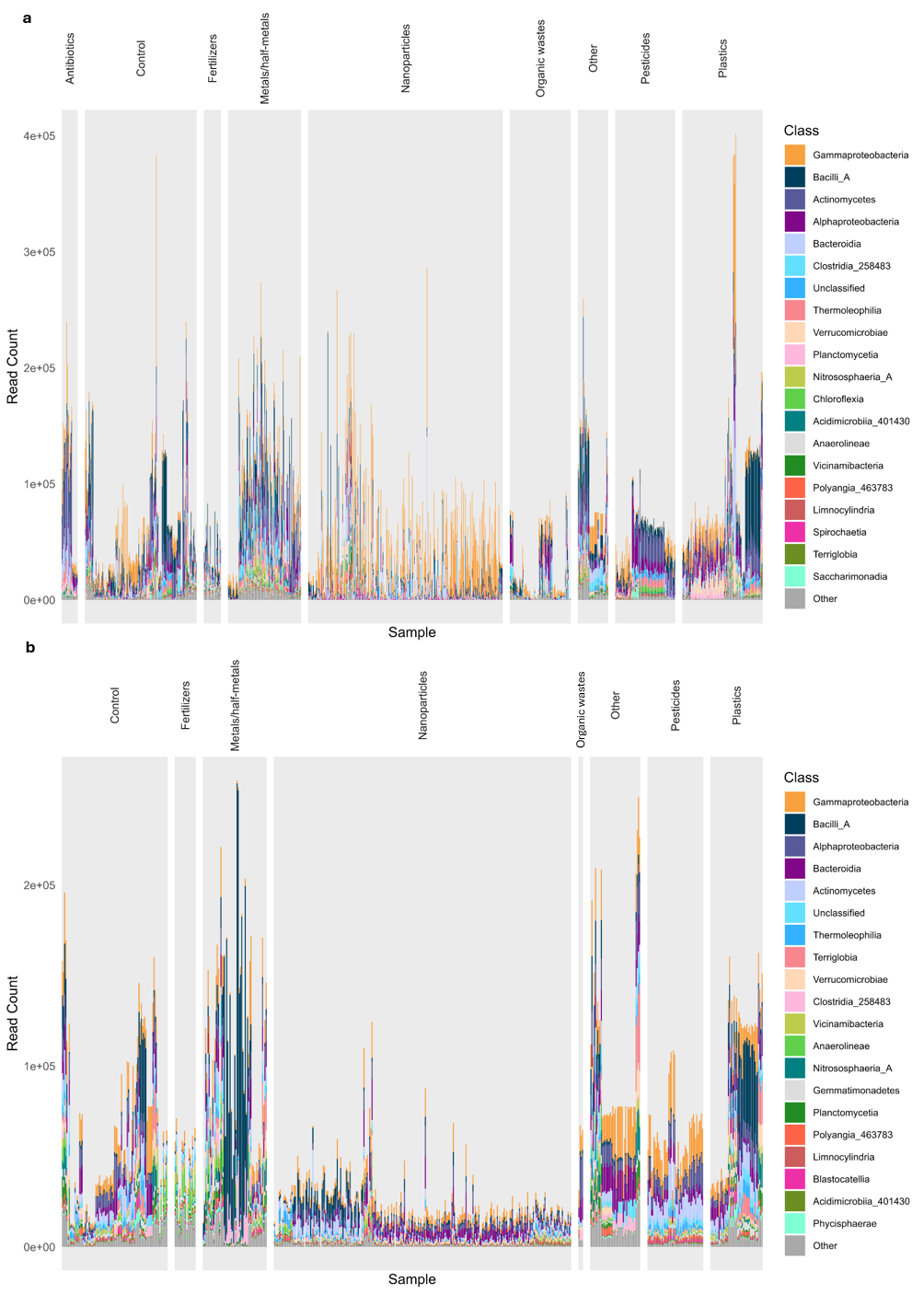
*Figure S8. Read counts for the different pollutant groups at the class level for A) internal (gut, cast) and B) external (skin, soil) samples with the top 20 most abundant taxa. Remaining taxa grouped together as ‘Other’. All unannotated taxa at the class level were grouped together as ’Unclassified’ to visualize the dominance of unclassified taxa in the samples.*

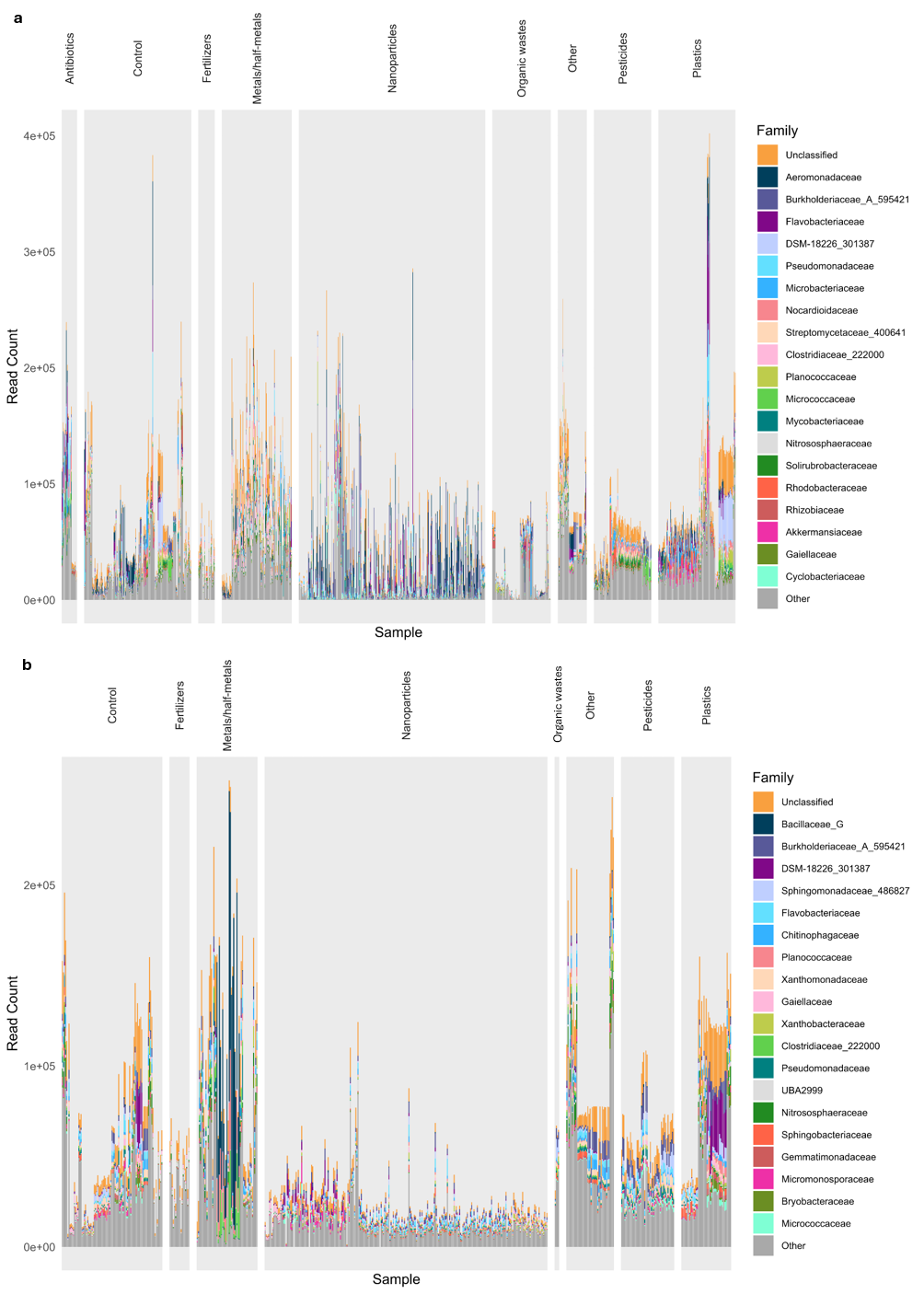
*Figure S9. Read counts for the different pollutant groups at the family level for A) internal (gut, cast) and B) external (skin, soil) samples with the top 20 most abundant taxa. Remaining taxa grouped together as ‘Other’. All unannotated taxa at the family level were grouped together as ’Unclassified’ to visualize the dominance of unclassified taxa in the samples.*

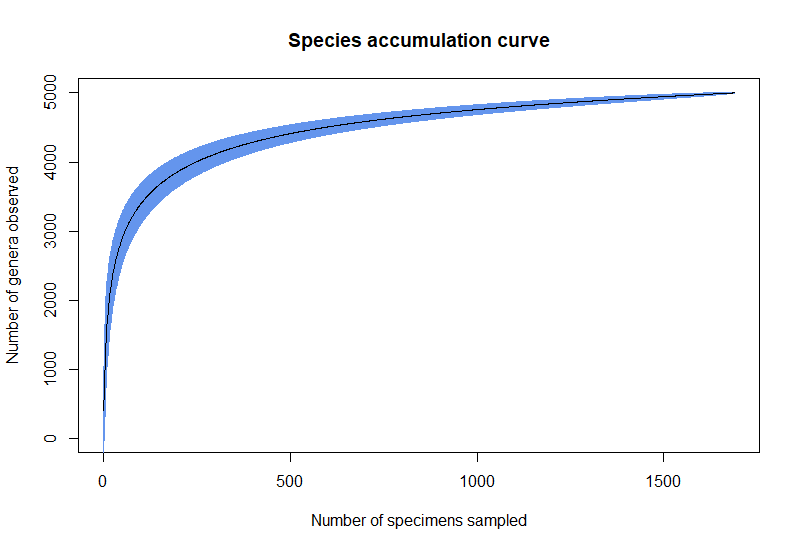
*Figure S10. Species accumulation curve for genus-level ASVs with 95 % confidence intervals. A plateau indicates good coverage.*
